# On-site metabarcoding analysis of environmental DNA samples

**DOI:** 10.64898/2026.03.27.714757

**Authors:** Quentin Mauvisseau, Isabelle Ewers, Isabella Blumeris, Sandra Irén Bongo, Luiz Filipe Brito de Oliveira, Bruno Gouvea, Aisha Carolina Cei, Karolina Ferreira Rodrigues, Jonata de Arruda Francisco, Even Sletteng Garvang, Victoria Marena do Rego Henriques, Sophia Hurtado Solano, Lone Kvalheim, Stephanie Kaylynne Lawrence, Bruna Ramalho Maciel, Hosea Isanda Masaki, Mahlatse Fortunate Mashaphu, Lindokuhle Masimula, Sankwetea Prudent Mokgokong, Emma Katrin Onshuus, Bianca Lima Paiva, Fatima Parker-Allie, Morne Du Plessis, Marco Puzicha, Orlando Gabriel Da Silva Solano Reis, Gift Speelman, Wilhelm Moritz Splitthof, Ana Carolina Stocco de Lima, Hanna Strindberg, Olea Smoge Sævik, Nicholas John Dillon Tafjord, Jorge Andres Varela Camelo, Lucia Josephine Yannuzzi, Trude Schmidt Øvregard, Rukaya Johaadien, Melania Rogowska, Dag Endresen, Cintia Oliveira Carvalho, Lisbeth Thorbek, Jarl Andreas Anmarkrud, Audun Schrøder-Nielsen, Adriana Lopes dos Santos, Micah Dunthorn, Hugo J. de Boer, Jonathan Ready

**Affiliations:** Natural History Museum, University of Oslo, Oslo, Norway; Department of Microbiology, Faculty of Science, Stellenbosch University, Stellenbosch, South Africa; Department of Biosciences, University of Oslo, Oslo, Norway; Group for Integrated Biological Investigation, Center for Advanced Studies of Biodiversity, Federal University of Pará, Belém, Brazil; Institute for Microbial Biotechnology and Metagenomics, University of the Western Cape, Bellville, South Africa; Marbio, Norwegian College of Fishery Science, Faculty of Biosciences, UiT-The Arctic University of Norway, Tromsø, Norway; Centre for Functional Biodiversity, Discipline of Biological Sciences, University of KwaZulu-Natal, Pietermaritzburg, South Africa; Agricultural Research Council, Pretoria, South Africa; University of Johannesburg, Johannesburg, South Africa; Unit of Environmental Sciences and Management, North-West University, Potchefstroom, South Africa; South African National Biodiversity Institute, Foundational Biodiversity Science, Cape Town, South Africa; University of the Free State, Bloemfontein, South Africa; Department of Chemistry, University of Oslo, Oslo, Norway; The School of Business, Innovation and Sustainability (FIH), Halmstad University, Halmstad, Sweden

**Keywords:** Environmental DNA, on-site analysis, Oxford Nanopore Technologies, MinION, BentoLab, protocol

## Abstract

Environmental DNA metabarcoding is a powerful monitoring tool for assessing aquatic biodiversity, as well as the sustainability and impacts of fisheries and aquaculture. However, conventional laboratory workflows remain time-consuming and dependent on dedicated infrastructures. Here, we present a field trial of a fully portable, off-grid eDNA metabarcoding pipeline that enables end-to-end analysis within a few days using compact equipment, including a BentoLab workstation and an Oxford Nanopore Technologies (ONT) MinION sequencer. The workflow was implemented during two international training courses in Norway and Brazil, where students and early career researchers collected environmental samples, extracted and amplified DNA, prepared DNA libraries, and sequenced on-site before performing bioinformatics and statistical analyses. In the case study detailed here, seven eDNA samples collected and analysed on-site in the Oslofjord allowed detection of 16 fish and elasmobranch species. Although overall diversity was lower than in earlier studies using Illumina-based sequencing, our protocol reliably detected key species and demonstrates that portable eDNA metabarcoding is feasible for rapid ecological assessment, surveillance of high-risk regions and/or deployment in remote or resourcelZllimited settings.

## Introduction

Anthropogenic activities, including habitat degradation, pollution, and climate change, are driving an ongoing mass extinction and severe global biodiversity loss (Ceballos et al., 2017). Aquatic ecosystems are especially affected, facing intensive pressures from capture fisheries and aquaculture (De Silva, 2012; Henriques et al., 2014), which, despite their importance for livelihoods and economies, impose substantial ecological costs (Béné et al., 2016; Campbell et al., 2021). These costs include ecological degradation, the spread of pathogens and parasites, and declines in wild fish populations (Henriques et al., 2014; Lafferty et al., 2015; Bayraktarov et al., 2016). The intensification of aquaculture further contributes to eutrophication, harmful algal blooms, and elevated disease-related mortality (Stentiford et al., 2012; Henriques et al., 2014; Liu et al., 2024). Training the next generation of scientists in fisheries and aquaculture is essential for monitoring and mitigating the environmental impacts of these industries while promoting sustainability. Environmental DNA (eDNA) metabarcoding offers an effective approach to assess ecosystem and fishery health (Gilbey et al., 2021; Ramírez-Amaro et al., 2022). While eDNA sample collection is relatively simple and can be carried out with minimal guidance (Agersnap & Thomsen, 2022; Burian et al., 2023; Clarke et al., 2023; Johnston et al., 2023), sample processing and data analysis require specialized training and facilities, are time-consuming, and often result in delayed outputs.

On-site processing of collected environmental DNA samples enables results within hours or days, allowing for near-real-time species detection and a more adaptive sampling design (Pomerantz et al., 2022; Patterson et al., 2025; Ip et al., 2026). Faster results and feedback allow researchers to rapidly adapt sampling intensity and location, adjust primer selection, and optimize laboratory processes or bioinformatic workflows. Extracting DNA shortly after sample collection also reduces its degradation and alleviates potential logistical issues associated with the transportation and preservation of eDNA samples in remote locations (Mauvisseau et al., 2021; Gygax et al., 2025; Ip et al., 2026). An established protocol for on-site eDNA metabarcoding further offers the possibility for performing regular ecosystem health assessments and biomonitoring activities, useful for aquaculture facilities that can better perform monitoring of pathogens, invasive species control, or manage harmful algal blooms.

Two international training courses were organized to provide students and early career professionals with hands-on experience on collecting, processing, and analyzing environmental DNA samples directly on site, outside of a dedicated laboratory or sequencing facilities. The course held in Bragança, Brazil, brought together 19 students from Brazil and Norway, while the one held in Drøbak, Norway, included 11 students and early-career researchers from Norway and South Africa. Through practical exercises using portable laboratory equipment and Oxford Nanopore Technologies (ONT) MinION sequencer, participants were trained in the development of sampling strategies, DNA extraction and amplification, basic bioinformatic and statistical analyses, and learned about research data management. Both courses emphasized the application of molecular tools to assess aquatic biodiversity, ecological health, and the sustainability of fisheries and aquaculture systems, thereby strengthening participants’ technical competence and fostering international research partnerships. In addition, by bringing together students from Brazil and Norway, these courses were designed as spaces of cultural and institutional exchange. Working side by side in mixed groups, students developed not only practical competence in metabarcoding, but also the ability to collaborate across languages, cultures, and institutional cultures. This intercultural dimension is crucial for the future of biodiversity monitoring in fisheries and aquaculture, which rely on transnational research networks and coordinated responses to ecological change.

Conducting all stages of the metabarcoding workflow outside a conventional laboratory setting presents unique logistical and technical challenges that can be exacerbated by environmental conditions (Zaiko et al., 2022). Extreme temperatures, strong winds, and rainfall can affect both sample integrity and personal performance, and careful planning is needed to mitigate this (Lin et al., 2025). Following the two field courses, we present here a reliable protocol that encompasses all steps of an off-grid metabarcoding analysis, from environmental sampling to end results. We used a portable on-site eDNA metabarcoding workflow encompassing collection of environmental DNA samples, DNA extraction, PCR amplification using two primer sets, library preparation, and sequencing on a compact and portable MinION sequencer. This was followed by off-grid bioinformatic processing of the raw sequencing output to generate a final OTU table and taxonomic assignment using a curated reference database. Custom R scripts were then used for quality control, perform basic analysis, and generate figures summarizing the findings. For brevity, we only show the analyses from the Norwegian site. The protocol and scripts provided here can be easily used and modified for teaching support or as a basis for an off-grid metabarcoding analysis in remote locations.

## Methods

### Environmental DNA sampling

Environmental DNA samples were collected following Carvalho et al. (2024a) and Mahlangu et al. (2025) in the Oslofjord at Drøbak (59°39’31.5“N 10°37’47.6“E) on the 3^rd^ and 4^th^ November 2025, and at the Aquarium of Drøbak (59°39’42.4“N 10°37’41.2“E) on the 4^th^ November 2025 (Table S1). In the Oslofjord, four eDNA samples were collected on the 3^rd^ November and two eDNA samples were collected on the 4^th^ November 2025 (Table. 1). At the Aquarium of Drøbak, one eDNA sample was collected on the 4^th^ November from the display tank harbouring the following fish and shark species: Thornback ray (*Raja clavata*), Pollock (*Pollachius pollachius*), Spotted catshark (*Scyliorhinus canicula*), Atlantic halibut (*Hippoglossus hippoglossus*), Ballan wrasse (*Labrus bergylta*), Turbot (*Psetta maxima*), Saithe (*Pollachius virens*), and Atlantic wolffish (*Anarhichas lupus*).

To minimize potential cross-contamination, unique sterile equipment and disposable gloves were used to collect and process each water sample. In both the Oslofjord and the Aquarium of Drøbak, surface water was collected using a sterile plastic bag (Whirl-Pak® 2041 mL Stand-Up Bag Merck®, Darmstadt, Germany), and each unique water sample (ranging from 750 mL to 2500 mL) was filtered using sterile 0.8 μm Whatman (Cytiva) mixed cellulose ester filters (22 mm diameter) and sterile 22 mm Swinnex (Merck Millipore) filter holders. Samples collected on 3^rd^ November were filtered using a sterile 50 mL Luer-Lock syringe BD Plastipak™, and using a Vampire Sampler (Bürkle GmbH) pump system on 4^th^ November. Filters collected on 3^rd^ November were immediately stored at −20 °C in 1.5 mL Eppendorf tubes until DNA extraction the following day, while samples collected on 4^th^ November were extracted immediately following water filtration.

### Environmental DNA extraction

Total DNA was extracted from each filter using the Qiagen DNeasy Blood & Tissue Kit (cat no. 69504) with slight modifications to the Quick-Start Protocol. Each filter was placed in a 1.5 mL tube, and double the volumes recommended by the manufacturer were added: ATL buffer (360 μl), proteinase K (40 μl), AL buffer (400 μl), and ethanol (400 μl). Lysis incubation with ATL buffer and proteinase K was performed in a thermal shaker at 56 °C and 500 rpm for 5 hours. The wash steps were performed as described in the protocol, and the elution was performed with 100 µl of 10 mM Tris-HCl per sample. Alongside the water filter samples, one extraction control (to which only reagents were added) was included. All centrifugation steps were performed in a BentoLab (Bento Bioworks Ltd, UK) centrifuge.

### PCR amplification

PCR amplifications targeting short fragments of the mitochondrial 12S *rRNA* were performed using the “Riaz” (12S-V5 Forward 5′-ACTGGGATTAGATACCCC-3′ and Reverse 5′-TAGAACAGGCTCCTCTAG-3′) and “Teleo2” (Forward 5′-AAACTCGTGCCAGCCACC-3′ and Reverse 5′-GGGTATCTAATCCCAGTTTG-3′) primer pairs (Riaz et al., 2011; Kelly et al., 2014; Taberlet et al., 2018; C. Carvalho et al., 2024, 2024) (Tables S2-S3). The Riaz primers amplified a 106 bp fragment, and the Teleo2 amplified an approximate (from 129 to 209 bp across taxa) 167 bp fragment. The primers listed above also had unique tags following the indexing strategies outlined in Fadrosh et al. (2014), and each eDNA sample was amplified with a unique tag combination in the forward and reverse primers (Table S4). Sequences of indexed primers with the associated samples can be found in Tables S2-3.

PCRs for each primer set were conducted in a final volume of 38 μL using 20 μL of 2x Accustart Toughmix II (QuantaBio), 1 μL of each indexed primer (5 µM each), 14 μL of nuclease-free water, and 2 μL of extracted eDNA. The PCR amplification parameters for both Riaz and Teleo2 primers included an initial denaturation at 94 °C for 2 min, followed by 35 cycles of 94 °C for 10 s, 55 °C for 20 s, and 72 °C for 15 s, with a final extension of 72 °C for 2 min. PCR reactions were conducted using a BentoLab (Bento Bioworks Ltd, UK).

All eDNA samples were analysed in four PCR replicates. The same indexed primer combination was used for a given sample, and PCR products were kept separate for subsequent indexing with different ONT barcodes (Figure 1). The extraction control sample was processed with two PCR replicates and later indexed with the ONT Native Barcodes NB01 and NB02. Additionally, two PCR controls made of ddH_2_O were also prepared (later indexed with ONT Native barcodes NB03 and NB04). Details of the indexed primer strategy used to analyze the samples are shown in Figure 1, and indexed primer sequences and index identifiers are provided in Tables S2-4.

**Figure 1.**
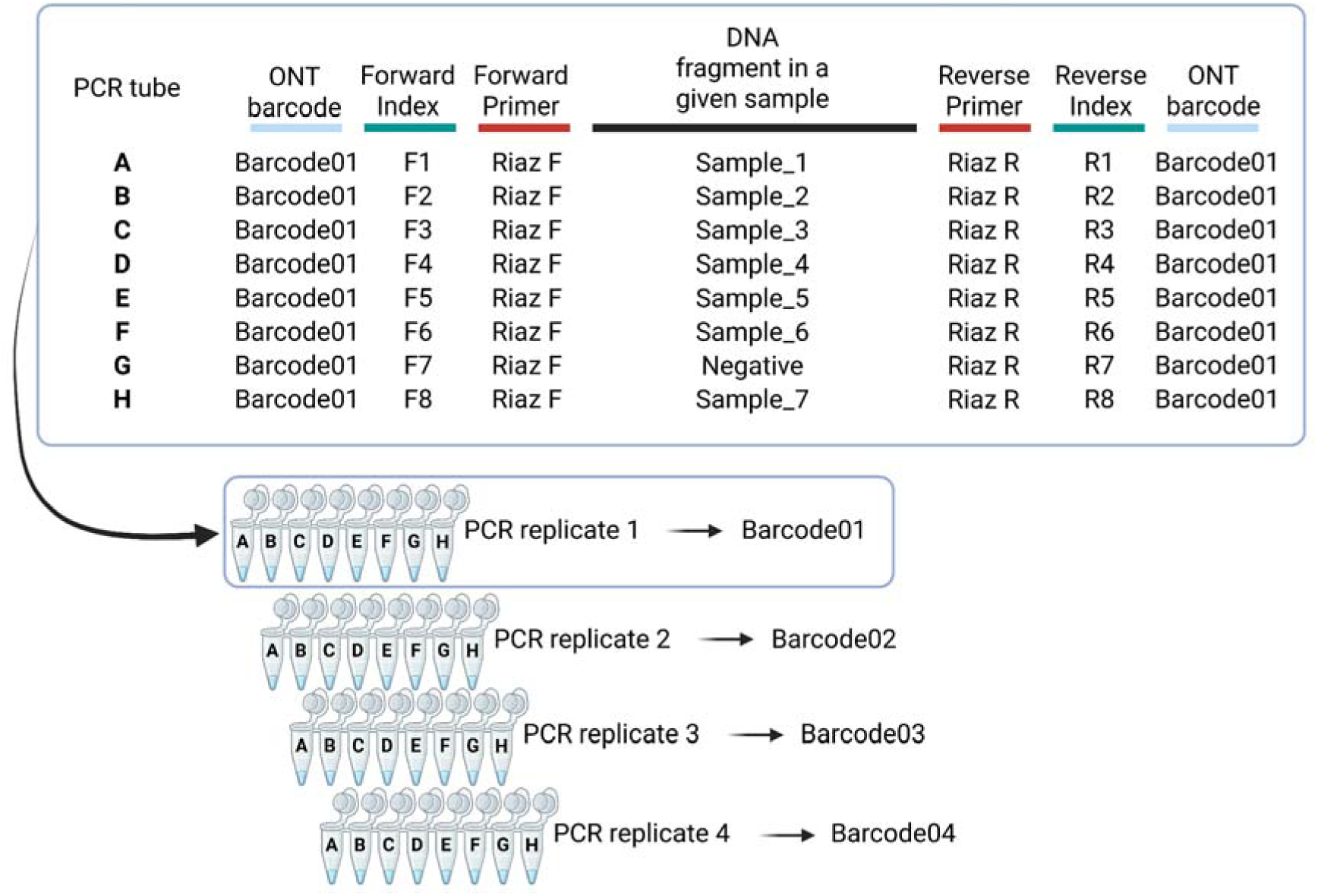
Workflow summarizing the strategy combining indexed primers and ONT barcodes to analyse the eDNA samples collected in this study with four PCR replicates. “Riaz F” refers to the forward primer, and “Riaz R” refers to the reverse primer. The workflow was identical for the Riaz and Teleo2 primer pairs.

**Figure 2.**
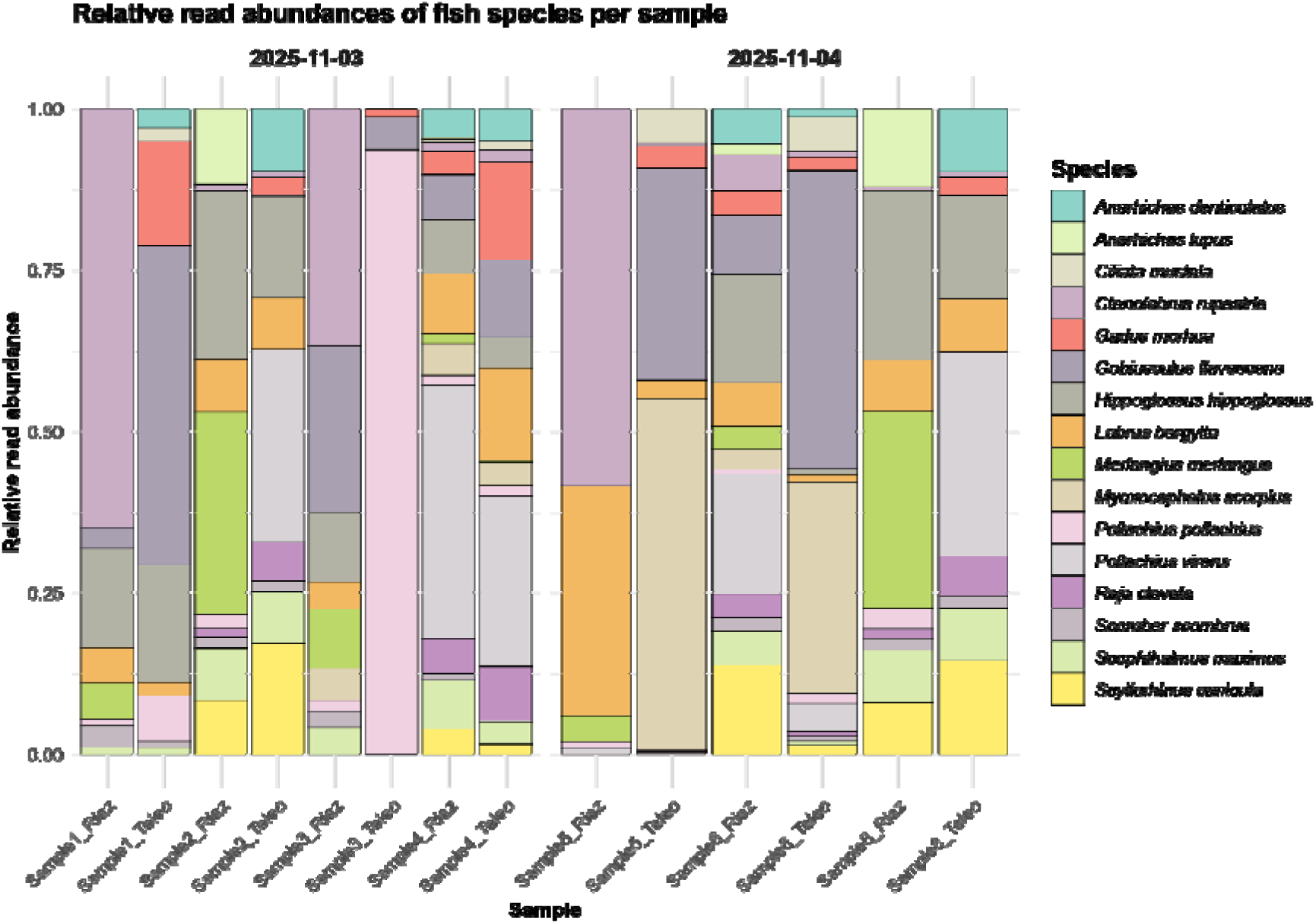
Relative read abundance of detected fish species in each sample from both sampling dates. Samples 1-6 were collected in the Oslofjord, while sample 8 was collected in Drøbak Aquarium.

### Library preparation, sequencing and bioinformatics

Library preparation was performed following a modified version of the SQK-NBD114-24 Minion protocol. All detailed steps and protocol can be found in the Supplementary Information section.

Sequencing was performed on a FLO-MIN114 flow cell using a SQK-NBD114-24 kit and the High-accuracy-v4.3.0 - 400bps basecall model with a minimum Q-score of 9 during 18 h 40 min. Samples from another project were also sequenced on the same flowcell using another ONT barcode. Each sample was demultiplexed using the cutadapt (version 5.2) software (Martin, 2011) based on the indexed primer sequences used for the PCR amplifications. Quality filtering was performed using VSEARCH v2.30.4_linux_x86_64 software (Rognes et al., 2016), removing short, low-quality reads, as well as sequences with ambiguous bases. Following this, all ONT barcode outputs were combined, sequences dereplicated, sorted by abundance, and filtered with a minimum occurrence threshold of 10 across the whole dataset using VSEARCH. Unique sequences were then clustered into Operational Taxonomic Units (OTUs) using a 97% identity threshold, chimeras were removed using the UCHIME3 algorithm as implemented in VSEARCH, and an OTU table was generated. Taxonomic assignment was carried out using both the SINTAX and usearch_global algorithms from VSEARCH, against the MIDORI2 database (Leray et al., 2022). The OTU table and taxonomic assignment were then used for downstream ecological analysis. Detailed commands for the bioinformatics processing can be found in Supplementary file 1.

### Biodiversity data management

Biodiversity research data was managed and published using FAIR data principles, following the DNA-derived data publishing guidelines from the Global Biodiversity Data Facility (GBIF) (Abarenkov et al., 2023).

### Statistical analysis

Data cleaning, manipulation, statistical analyses, and visualization were conducted in R version 4.5.1 (R Foundation for Statistical Computing, 2025). Using the packages readr v. 2.1.5 (Wickham et al., 2015), dplyr v. 1.1.4 (Wickham, François, et al., 2014), and stringr v. 1.5.2 (Wickham, 2009), the taxonomic assignment file generated using the SINTAX algorithm in VSEARCH was cleaned by parsing the rank and similarity score of each taxonomic assignment entry to remove rank prefixes and taxIDs, and combined with the raw OTU table. A data frame containing a summary of the metadata available for each sample was additionally created to be used for diversity analyses and data comparisons (Table S1). Then, the OTU table was filtered and cleaned to remove non-target taxa, and potential contaminants or sequencing errors. As fish and elasmobranchs were the primary focus of this study, only OTUs assigned to the classes “Actinopterygii” or “Chondrichthyes” within the phylum “Chordata” were retained. To correct for potential contamination, the maximum number of reads observed across the four replicates of the negative control for each primer set was subtracted from all corresponding samples. This procedure was applied to each unique OTU before removing those with a resulting null read count across all samples, as well as the negative control from the OTU table. The remaining OTUs were further filtered to retain only those assigned to the family level or lower with a taxonomic identity score of ≥=0.99. Finally, singletons (i.e., OTUs with one read across all replicates and samples) were removed. To obtain the final OTU table for the statistical analyses, each species-level taxonomic assignment was validated by cross-referencing species occurrence records in the Oslofjord area from Artsdatabanken and the Global Biodiversity Information Facility (GBIF) database (*GBIF Derived Dataset*, 2025). OTUs assigned to species not reported in this area were checked against the NCBI database using BLAST, and corrected assignments were incorporated into the filtered table. OTUs assigned to the same species were then pooled, as well as PCR replicates of any unique sample for each primer set.

Analyses and comparisons of fish diversity were performed using the packages dplyr v. 1.1.4 (Wickham et al., 2014), tidyr v. 1.3.1 (Wickham et al., 2014), and vegan v.2.7-1 (Oksanen et al., 2001). Visualizations were generated with ggplot2 v. 3.5.2 (Wickham, 2016), patchwork v. 1.3.2 (Pedersen, 2019), and RColorBrewer v. 1.1-3 (Neuwirth, 2002). Species accumulation curves were plotted for each primer set to estimate sampling effort, followed by species richness, relative read abundance, and Shannon diversity calculation per sample. Diversity comparisons by primer sets and sampling date were visualized using boxplots. To assess community (dis)similarities between primers and sampling dates, a PERMANOVA test was performed on a presence-absence version of the data, and beta-diversity was visualized using NMDS plots. Stacked bar plots were used to visualize species-level relative abundance and composition between samples. Venn diagrams were generated to compare species detected by the two primer sets, between field samples and positive controls, and across sampling dates. Species accumulation curves, Shannon diversity boxplots, NMDS plot, and Venn diagrams are presented in the supplementary information section (Figures S1-S4), while the relative abundance barplots are shown in the Results and Discussion section. Due to the different output format generated by the usearch_global algorithm in VSEARCH for the taxonomic assignment step, the R script for importing and processing the taxonomic assignment table was adapted for the data set “hits.blast6”. The modifications were implemented in section 1.3 of the R-script used for the data processing, analysis, and visualization. This script, along with the raw sequencing data,can be found in the following Zenodo repository (DOI:10.5281/zenodo.18222631).

## Results and Discussion

The MinION sequencing run lasted 18 h 40 min, generating 13.43M reads and approximately 8.04 Gb of called bases. Amongst them, 3.0M, 2.7M, 2.3M, and 3.1M reads passed basecalling quality thresholds in Barcode01, Barcode02, Barcode03, and Barcode04 respectively. After metabarcoding processing and filtering, a total of 3,093,315 raw reads assigned to 1,053 unique OTUs were obtained. From the negative controls, a maximum number of 110 reads in a single OTU were obtained using the Riaz primer set, and 554 reads in a single OTU were obtained by the Teleo2 primer set. Those reads belonged to the same OTU assigned to *Pollachius virens*, and were subtracted from all other samples. Subsequent quality filtering, removal of potential contaminants using negative controls, retention of OTUs assigned to fish and elasmobranch, manual correction of taxonomic misidentifications, and pooling of OTUs assigned to the same species, yielded 205,381 reads representing 16 fish and elasmobranch species.

The taxonomic assignments of the 27 unique fish and elasmobranch OTUs recovered from the eDNA samples were corrected for six OTUs prior to combining OTUs assigned to the same species. These six unique OTUs, initially assigned to five different species using the MIDORI2 reference database, were identified as false positives due to the absence of occurrence records for these species in the Oslofjord area according to Artsdatabanken and GBIF records (*GBIF Derived Dataset*, 2025). BLAST comparisons of the corresponding DNA sequences against the NCBI database revealed higher similarity scores ≥ 99% to four closely related species, compared to the original MIDORI2 assignments (77-96%) (Leray et al., 2022). Specifically, OTUs identified as *Dipturus batis* (blue skate) and *Raja asterias* (starry skate) were reassigned to *Raja clavata* (thornback ray), both *Eopsetta grigorjewi* (shotted halibut) OTUs were corrected to *Hippoglossus hippoglossus* (Atlantic halibut), *Gadus chalcogrammus* (Alaska pollock) to *Gadus morhua* (Atlantic cod), and *Myoxocephalus aenaeus* (grubby) to *Myoxocephalus scorpius* (shorthorn sculpin). All four corrected species are known to occur in the Oslofjord area (Carvalho et al., 2024a; Kvalheim et al., 2024; *GBIF Derived Dataset*, 2025; Mahlangu et al., 2025), underscoring the importance of incorporating local species occurrence records in the taxonomic validation of eDNA metabarcoding results. The false-positive assignments likely resulted from the short amplicon lengths of the Riaz and Teleo2 primer sets (127-188 bp), which may not provide sufficient resolution for accurate species delimitation based on the 12S marker (Burian et al., 2021, 2023). Another contributing factor could be biases introduced by the SINTAX algorithm in VSEARCH, which relies on k-mer similarity patterns and automatically assigns the most similar taxon available in the available reference database (Edgar, 2016, Carvalho et al., 2024a).

No significant differences were observed in the community composition based on presence-absence data between the two primer sets, with both Riaz and Teleo2 detecting the same 16 species. Similarly, no significant differences were found between the fish and elasmobranch community detected across the two sampling dates. All eight fish species identified in the aquarium samples (positive control) using eDNA metabarcoding were also detected in the environmental samples from the Oslofjord. Additionally, eight more fish species than those recorded as physically present in the aquarium tank were detected in the positive control samples, likely reflecting the continuous intake of seawater from the Oslofjord.

When comparing our results with previous eDNA surveys conducted in the Oslofjord during the same season, we identified five species in our dataset that had not been detected in any of the other studies (Carvalho et al., 2024a; Kvalheim et al., 2024; Mahlangu et al., 2025). Using the Riaz and Teleo2 primer sets, we detected *Anarhichas denticulatus* (northern wolffish), *Anarhichas lupus* (Atlantic wolffish), *Hippoglossus hippoglossus* (Atlantic halibut), *Raja clavata* (thornback ray), and *Scyliorhinus canicula* (small-spotted catshark) in both the aquarium (positive control) and environmental samples from the Oslofjord. The recovery of these species compared to previous eDNA surveys of the same region and season likely reflects differences in primer choices. In contrast to our approach, Carvalho et al. (2024a), Kvalheim et al. (2024), and Mahlangu et al. (2025) used the MiFish (Miya et al., 2015; Sales et al., 2019) and Elas02 (Taberlet et al., 2018) primer sets, which did not detect these five species. This highlights the significant influence of the primer set selection in the detection of specific fish and elasmobranch taxa in eDNA metabarcoding studies and underscores the need to account for potential primer biases when interpreting biodiversity assessment derived from such data (Bylemans et al., 2018; Schenekar et al., 2020; Zhang et al., 2020; Kumar et al., 2022; Burian et al., 2023). To mitigate potential issues associated with specific primer set or reference database coverage, a multi-marker approach should be considered when possible (Stefanni et al., 2018; Alexander et al., 2019; Cordier et al., 2019; Wang et al., 2021; Brys et al., 2023; Burian et al., 2023; Zhu & Iwasaki, 2023; Carvalho et al., 2024a; Espinosa Prieto et al., 2024).

However, when comparing with previous eDNA surveys from the Oslofjord area, our study recovered only a fraction of the reported fish diversity. While earlier surveys detected 51 to 65 fish species (Carvalho et al., 2024a; Kvalheim et al., 2024; Mahlangu et al., 2025), our results revealed a total of 16 species. This difference likely reflects the sampling design applied in our study, which involved a limited number of samples collected at a single sampling location over a short time period. Indeed, such restricted spatio-temporal coverage combined with a low volume of water filtered and absence of natural replication are known factors leading to false negative results in eDNA surveys (Ficetola et al., 2015; Weigand & Macher, 2018; Beentjes et al., 2019; Mauvisseau et al., 2019; Shirazi et al., 2021; Stauffer et al., 2021; Anmarkrud et al., 2025). Additionally, previous studies relied on sequencing through the Illumina MiSeq platform, whereas our study used the Oxford Nanopore MinION platform, the latter being known for its higher sequencing error rate (Baloğlu et al., 2021; Delahaye & Nicolas, 2021; Egeter et al., 2022; Doorenspleet et al., 2025; Gygax et al., 2025; Patterson et al., 2025). To mitigate this, we applied stringent bioinformatic filtering thresholds, which may have inadvertently led to over-filtering of raw reads and the loss of true detection, further contributing to the lower level of recovered diversity.

In field situations away from clean labs, we were able to complete the entire environmental DNA metabarcoding workflow, from sampling, DNA extraction, amplification, library preparation, sequencing, and data processing on-site and within five days. The successful completion of all steps outside a dedicated laboratory and without access to conventional sequencing facilities, all while using portable equipment, demonstrates that data can be collected and analyzed in remote locations with limited resources. Portable laboratory tools such as the BentoLab system, a DIYNAFLUOR device (a 3D printed DNA fluorometer for self-assembly, (Anderson et al., 2024)), and a MinION sequencer, together with a bioinformatics pipeline designed to run on a standard laptop without internet access, substantially increase the feasibility of this workflow under unpredictable or otherwise unfavorable conditions. Despite the potential limitations highlighted earlier, participants were able to complete the entire metabarcoding workflow within the duration of the courses. When all critical aspects are carefully accounted for, the on-site metabarcoding workflow described here has the potential to address a wide range of research questions within a short timeframe.

## Conclusion

Here we provided a practical and do-able approach to performing eDNA metabarcoding in the field when away from clean labs and away from high-capacity computers. This approach can easily be modified to work on other primer pairs and gene regions. The bioinformatics codes we provided are portable across laptop platforms. These codes can also easily be modified to meet specific needs and can integrate additional bioinformatics steps. Such a practical approach is beneficial for researchers as it enables rapid on-site diagnostics and environmental assessments, with the potential to provide critical information for decision-makers in environmental agencies or the fishery sector. The open-access data and protocols provided to reproduce the entire workflow further support its use in other teaching courses and as a basis for investigating diverse research questions.

## Acknowledgements

We would like to thank the staff of Drøbak Aquarium, Norway, for allowing us to collect eDNA samples in their facility. We thank Grete Sørnes from Drøbak Marine Station and the technicians from the DNA lab at the Natural History Museum at the University of Oslo for their support and assistance when planning the two different courses.

## Author Contribution

Design: QM, IE, MD, HdB, DE, MdP, FpA, JSR; Bioinformatics: QM and IE; Data curation: QM and IE; Statistical analysis: QM and IE; Original draft: QM and IE; Fieldwork, Laboratory processing, Review and editing: all co-authors; Funding acquisition: QM, HdB, MD, DE, MdP, FpA, JSR; Supervision: QM, HdB, MD, MdP, FpA, JSR.

## Data availability statement

Raw sequencing data, OTU table, taxonomic assignments, and R scripts can be found in the following Zenodo repository (DOI:10.5281/zenodo.18222631). Additional information linked to sites sampled, primer sequences, and indexes can be found in Supplementary information.

## Supplementary information

Supplementary file 1.

### Library preparation

Library preparation was performed following a modified version of the SQK-NBD114-24 Minion protocol. First, all indexed amplicons corresponding to a given PCR replicate (i.e. PCR replicates 1, 2, 3 or 4) were pooled into a single tube. For example, the 16 PCR products corresponding to the amplification of the eDNA samples and negative control with the Riaz and Teleo2 primer set were pooled together (8 samples with Riaz and 8 with Teleo2, see Figure 1). This resulted in four pooled samples, one for each of the four PCR replicates for Riaz and Teleo2 primer sets. The four pooled samples were kept separate, then each pool was cleaned with Ampure XP in a 1:1 ratio and eluted in 200 µl before concentration was measured using a Qubit. Approximately 200 fmol was used as input in each End-prep reaction consisting of a total of 16 μL of diluted DCS, 14 μL of Ultra II End-prep rxn buffer, and 6 μL of Ultra II End-prep Enzyme (all maintained at 4 °C prior to use) were added to each pooled sample. The pooled samples were then incubated at 20 °C for 5 min, followed by 65 °C for 5 min,to inactivate remaining enzymes.

The four pools were purified with 1X AMpure XP beads before elution in 6 µl dH_2_O. To do so, freshly diluted 70% ethanol was prepared, and 550 µL was added in each pool. A 1:1 ratio of Ampure XP beads was then added to each amplicon pool, and incubated at room temperature for 5 min with gentle mixing. Tubes were placed on a magnetic rack for 2 min until the solution cleared, and the supernatant was carefully pipetted off without disturbing the bead pellet. 250 µL of 70% ethanol was added while the tubes were still on the magnet for a washing step, then the ethanol was removed after 1 min. This ethanol washing step was repeated twice. Then, the tubes were briefly centrifuged and put back on the magnetic rack, and any remaining ethanol was removed. Tubes were then left open for 5 min to air dry the pellets. Subsequently, 35 µL of nuclease free water was added to each tube while on the magnet. The tubes were then closed and removed from the magnet, gently flicked to resuspend the beads in the water, and left at room temperature for 1 min before being placed back on the magnet for 2 min until the solution cleared. This was done to resuspend the DNA into the water. 6 µL of the clear solution containing the DNA was then pipetted into new tubes.

Following this, different MinION Native Barcode Adapters were ligated to the four separate pools. Barcode01 was ligated to the pool corresponding to the first PCR replicate (Figure 1), Barcode02 was ligated to the pool corresponding to the second PCR replicate, Barcode03 was ligated to the pool corresponding to the third PCR replicate, and Barcode04 was ligated to the pool corresponding to the fourth PCR replicate. A total of 3.75 µL of the end-prepped DNA (resuspended as described above) was combined with 1.25 µL of the corresponding Native Barcode Adapter and 5 µL of Blunt/TA Ligase Master Mix for a final volume of 10 µL, and incubated at 20 °C for 20 min. Following incubation, 2 µL of EDTA was added to each pool followed by mixing to inactivate the ligase.

Subsequently, all sample pools indexed with their different Native Barcode Adapters were combined into a single tube, and purified 1X Ampure XP beads were added to the final pool and incubated at room temperature for 5 min with gentle mixing. The tube was then placed on a magnetic rack for 2 min until the solution cleared, and the supernatant was carefully pipetted off without disturbing the bead pellet. 250 µL of 70% ethanol was added while the beads were still on the magnet for a washing step, then the ethanol was removed after 1 min. This ethanol washing step was repeated twice.The tube was briefly centrifuged and put back on the magnetic rack, and any remaining ethanol was removed. Tubes were then left open for 5 min to air dry the pellets, then, 32 µL of nuclease free water was added to the tube while on the magnet. The tube was then closed and removed from the magnet, gently flicked to resuspend the beads in the water, and left at room temperature for 1 min before being placed back on the magnet for 2 min until the solution cleared. This was done to resuspend the DNA into the water. 1 µL of the purified elute was quantified using a QuBit.

Following this, Native Adapter ligation was performed. 30 µL of the pooled barcoded sample was combined with 5 µL of the MinION Native Adapter, 10 µL of the Quick ligase reaction buffer, and 5 µL of the Quick T4 ligase for a final volume of 50 µL, and incubated at 20 °C for 20 min. Immediately after incubation, the library was purified with Ampure XP: a 1:1 ratio of Ampure beads was added to the pool, and incubated at room temperature for 5 min with gentle mixing. The tube was then placed on a magnetic rack for 2 min until the solution cleared, and the supernatant was carefully pipetted off without disturbing the bead pellet. 125 µL of SFB was added while the beads were still on the magnet for a washing step, then the SFB was removed after 1 min. This SFB washing step was repeated twice.The tube was briefly centrifuged and put back on the magnetic rack, and any remaining SFB was removed. Tubes were then left open for 5 min to air dry the pellets, then, 15 µL of elution buffer was added to the tube while on the magnet. The tube was then closed and removed from the magnet, gently flicked to resuspend the beads in the elution buffer, and left at room temperature for 1 min before being placed on the magnet again for 2 min until the solution cleared. 1 µL of the cleaned elute was quantified using a QuBit.

Finally, the priming mix consisting of 1170 µL of FCF, 12,5 µL of BSA (20 mg/mL), and 30 µL of FCT was used to prime the flow cell. First, the flow cell was placed in a MK1D MinION sequencer. 15 µL of the buffer was dialed up from the priming port to remove potential air bubbles in the channel. Then, using new pipette tips, 800 µL of the priming mix was loaded into the flow cell via the priming port. The priming port cover was then left open for 5 min, after which 200 µL of the priming mix was slowly loaded in the SpotON port. Immediately following priming, a final 75 µL volume consisting of 12 µL of the final library mixed with 37.5 µL of SB and 25.5 µL of LIB was loaded dropwise onto the SpotON port. The port was then closed, the light cover placed, and the MK1D sequencer was connected to the laptop, and the sequencing run was initiated through the MinKNOW software interface.

### Bioinformatics processing

The metabarcoding analysis was performed on the reads present in the “fastq_pass” folder, where the folders containing the results of each ONT barcode can be found. Some of the bioinformatic processes were performed in each of these barcode’s folders, and the remaining of the analysis were conducted in another dedicated folder. Prior to analysis, a dedicated “mapping file” summarizing the sample IDs, PCR replicate numbers, and sequences of the indexed forward and reverse primers was placed in each of the “barcode” folder alongside the fastq.gz files. Raw sequencing data and mapping files corresponding to each barcode folder can be found in (DOI:10.5281/zenodo.18222631). In the “fastq_pass” folder the following commands were ran:

**Step 1** - We created a new folder to pool and combine all the PCR replicates together for processing.

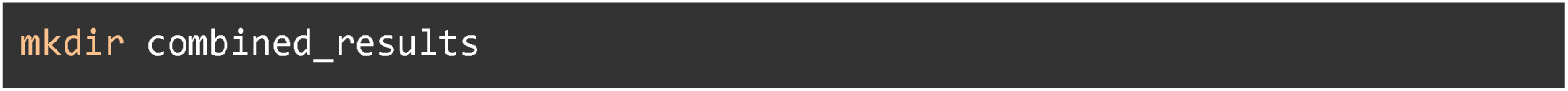

The steps 2 - 4 detailed below were run for all barcode folders. The example provided was run in the barcode01 folder.

**Step 2** - We unzipped and merged all the fastq.gz files for processing.

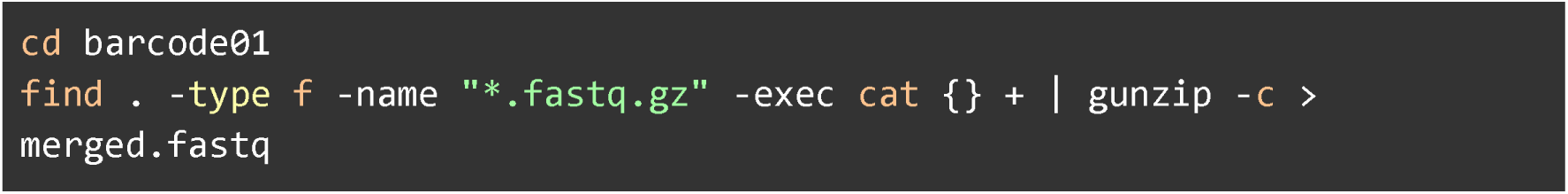

**Step 3** - We demultiplexed all samples corresponding to the first PCR replicate using the mapping file including the sample ID, PCR replicate number, and indexed primer sequences.

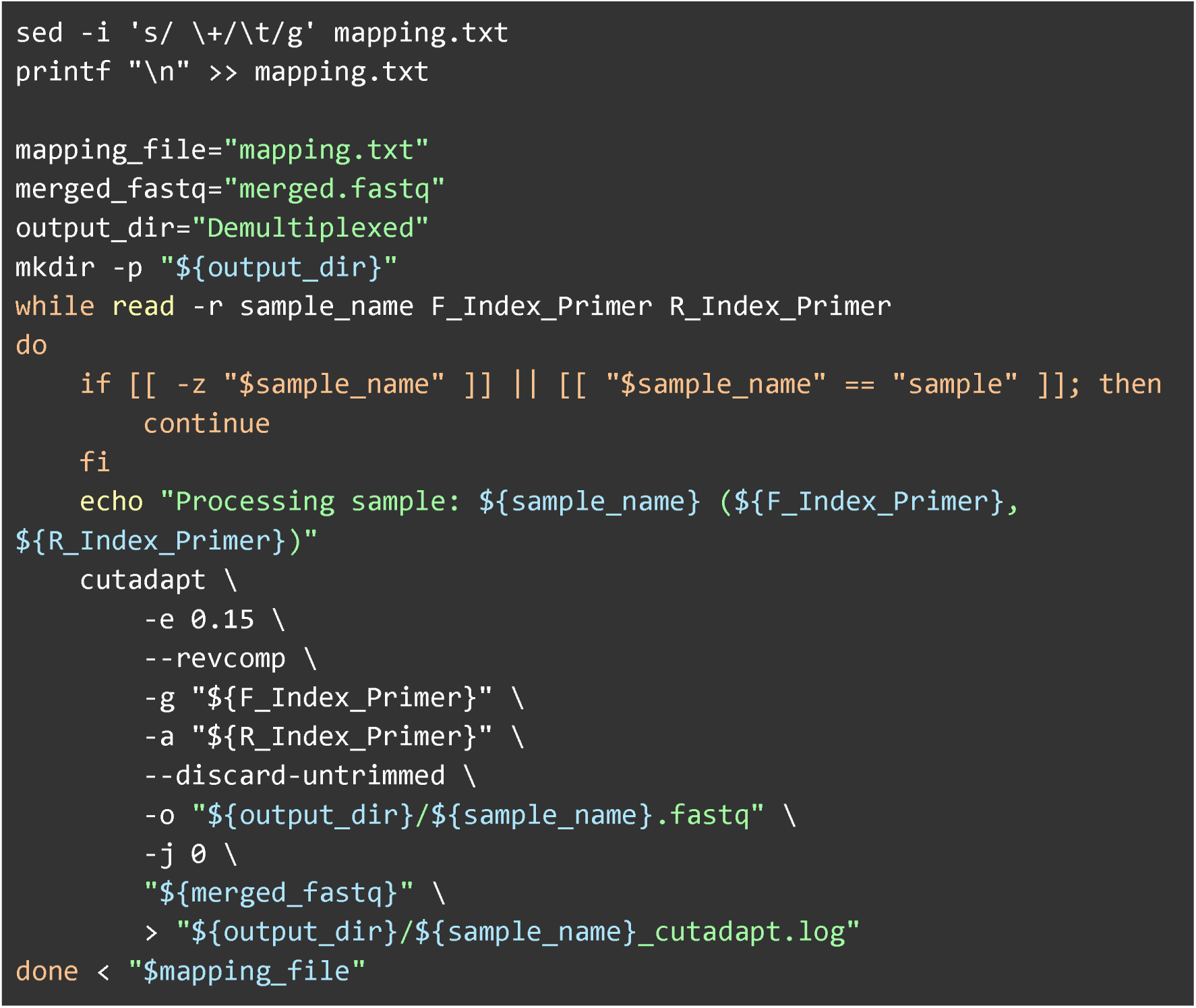

**Step 4** - We used VSEARCH for quality control, and removed reads with >5 expected errors, shorter than 50 bp, and containing any unknown base.

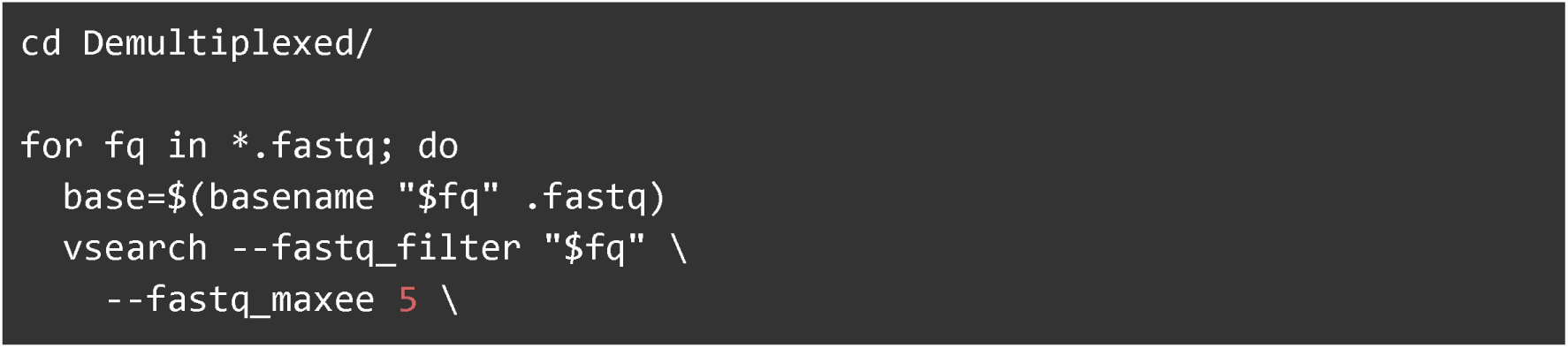
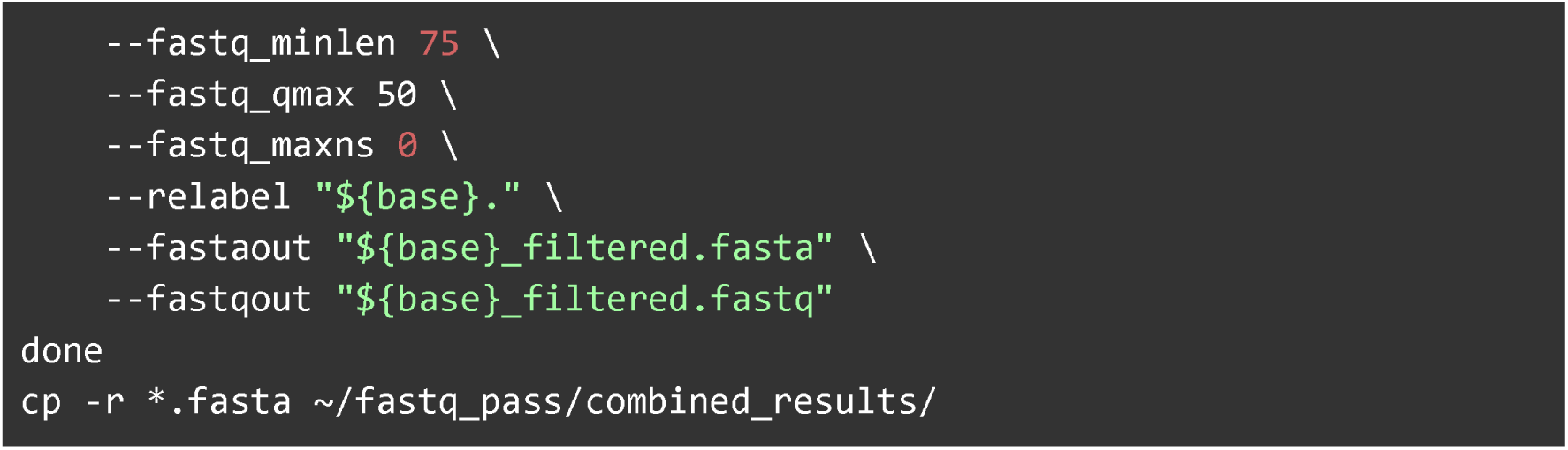

The Steps 2-4 described above should be run in each barcode folder. All their outputs were copied into the “combined_results” folder for further processing.

**Step 5** - We combined all filtered fasta files, dereplicated the sequences using VSEARCH, sorted them by size, and removed unique sequences with less than 10 occurrences in the whole dataset.

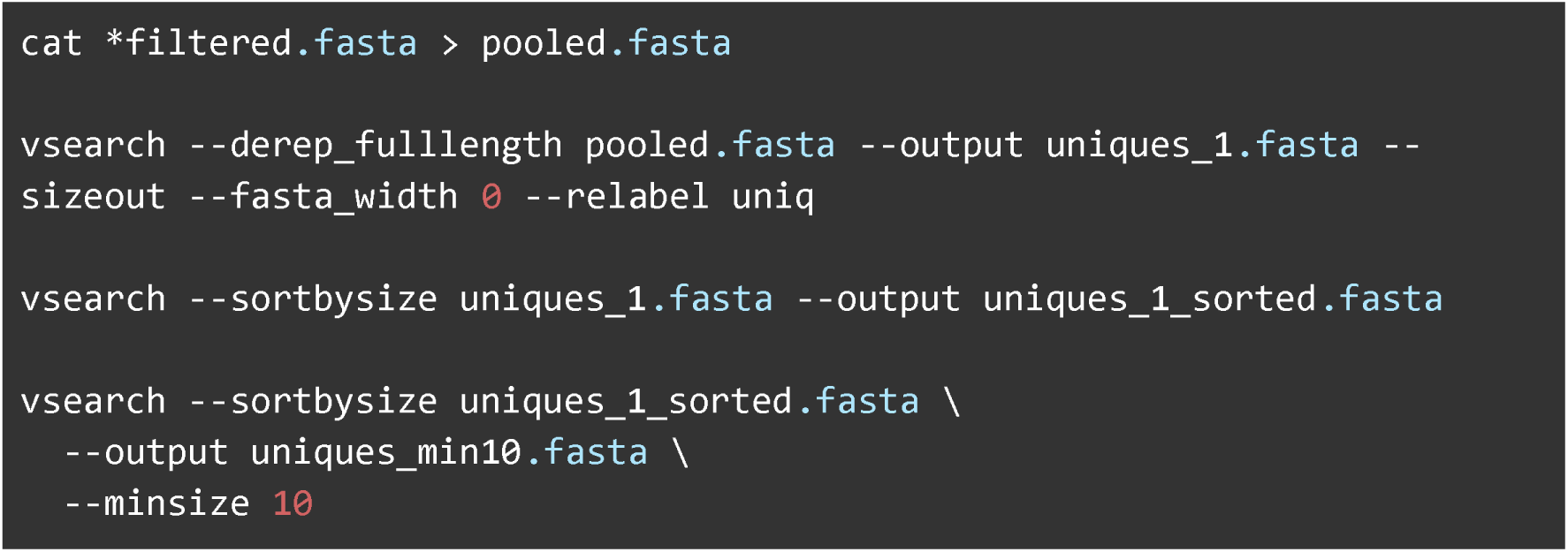

**Step 6** - We denoised the dataset using 0.97% identity, removed chimeras, and generated an OTU (Operational Taxonomic Unit) table.

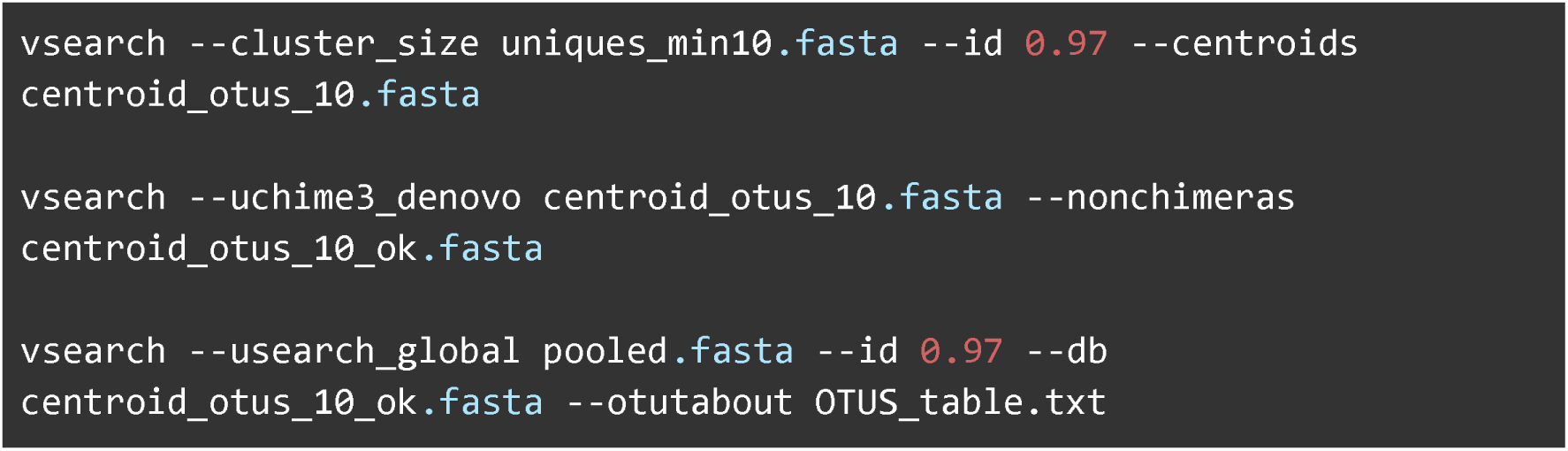

**Step 7** - We performed the taxonomic assignment of the generated OTUs using VSEARCH. The MIDORI2 database (release 269) was downloaded from https://www.reference-midori.info/download/Databases/GenBank269_2025-12-09/SINTAX_sp/uniq/MIDORI2_UNIQ_NUC_SP_GB269_srRNA_SINTAX.fasta.gz, and taxonomic assignment was conducted using the SINTAX algorithm in VSEARCH. First, the <u>fasta.gz</u> file of the MIDORI2 database was downloaded to the “combined results” folder. The following code was then used to convert the database to the VSEARCH format, and the taxonomic assignment was run with a 0.95 cutoff using both SINTAX and usearch_global algorithms. Data processing, analysis and visualization were later done using the taxonomic assignment obtained using SINTAX. R-scripts necessary to run similar analysis on the taxonomic assignment obtained using the usearch_global algorithm can be found in the supplementary information section.

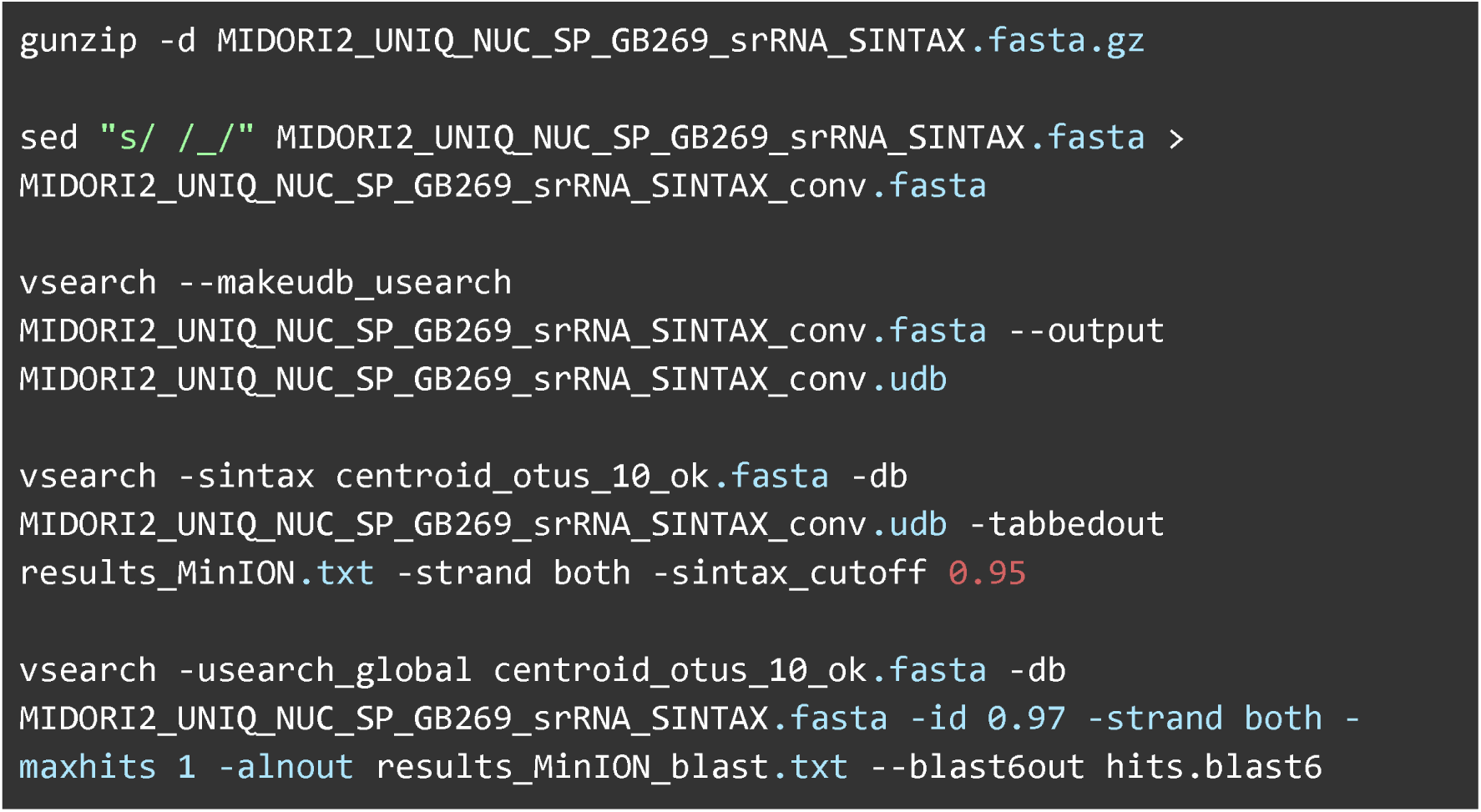

**Table S1.** Overview of the volume, sampling date and location of the samples collected and analysed in this study.

| Sample ID | Volume filtered | Date | GPS coordinates* |
| --- | --- | --- | --- |
| Sample_1 | 750 mL | 03/11/2025 | 59°39'31.5"N<br>10°37'47.6"E |
| Sample_2 | 750 mL | 03/11/2025 | 59°39'31.5"N<br>10°37'47.6"E |
| Sample_3 | 750 mL | 03/11/2025 | 59°39'31.5"N<br>10°37'47.6"E |
| Sample_4 | 750 mL | 03/11/2025 | 59°39'31.5"N<br>10°37'47.6"E |
| Sample_5 | 800 mL | 04/11/2025 | 59°39'31.5"N<br>10°37'47.6"E |
| Sample_6 | 1000 mL | 04/11/2025 | 59°39'31.5"N<br>10°37'47.6"E |
| Sample_7 | - | 04/11/2025 | Extraction control |
| Sample_8 | 2500 mL | 04/11/2025 | 59°39'42.4"N<br>10°37'41.2"E |

\* Samples 1-6 were collected in the Oslofjord (59°39'31.5"N 10°37'47.6"E) while sample 8 was collected at the Drøbak Aquarium (59°39'42.4"N 10°37'41.2"E).

**Table S2.**
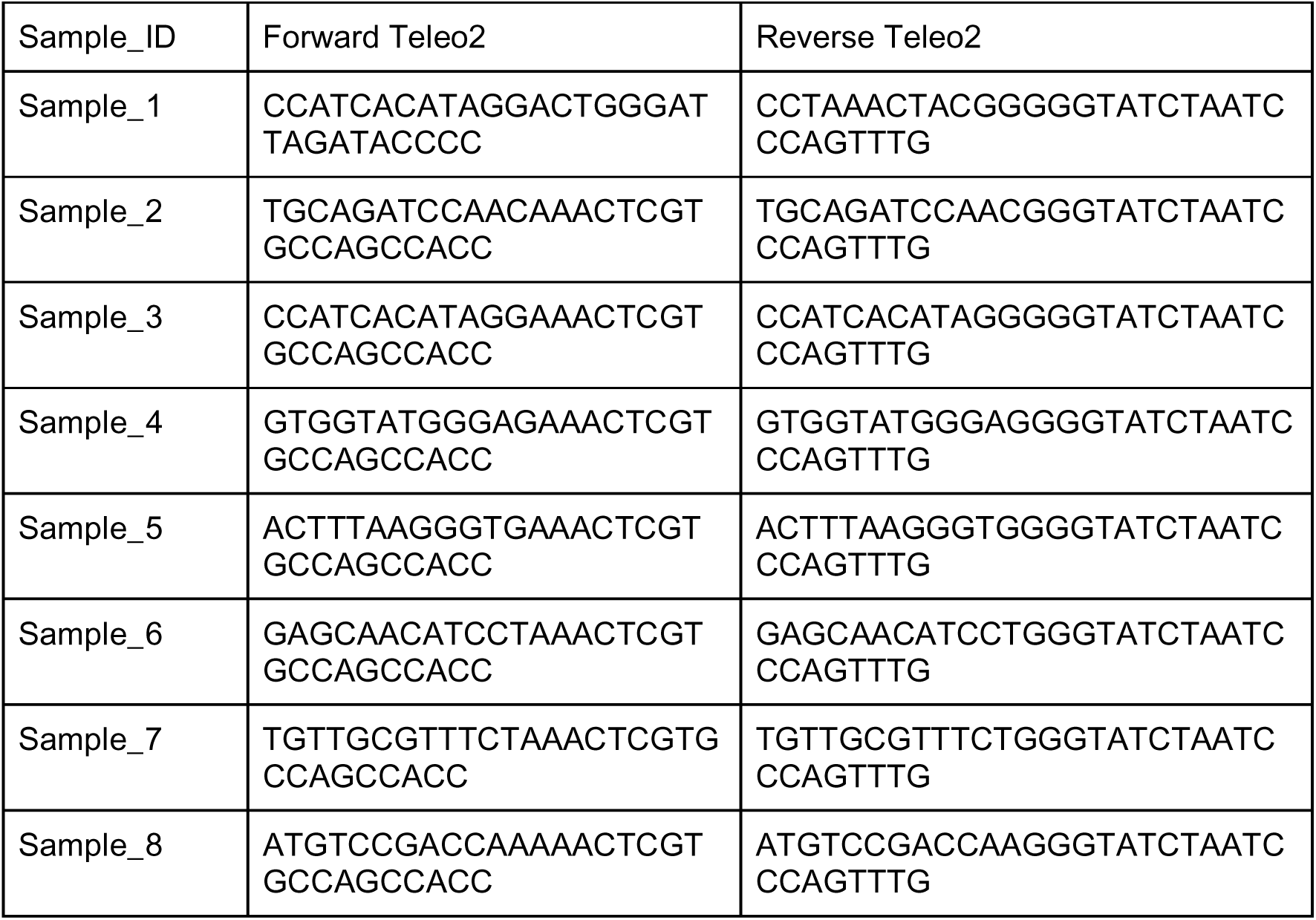
Sequences of the indexed forward and reverse Teleo2 primers in the 5’-3’ direction.

**Table S3.**
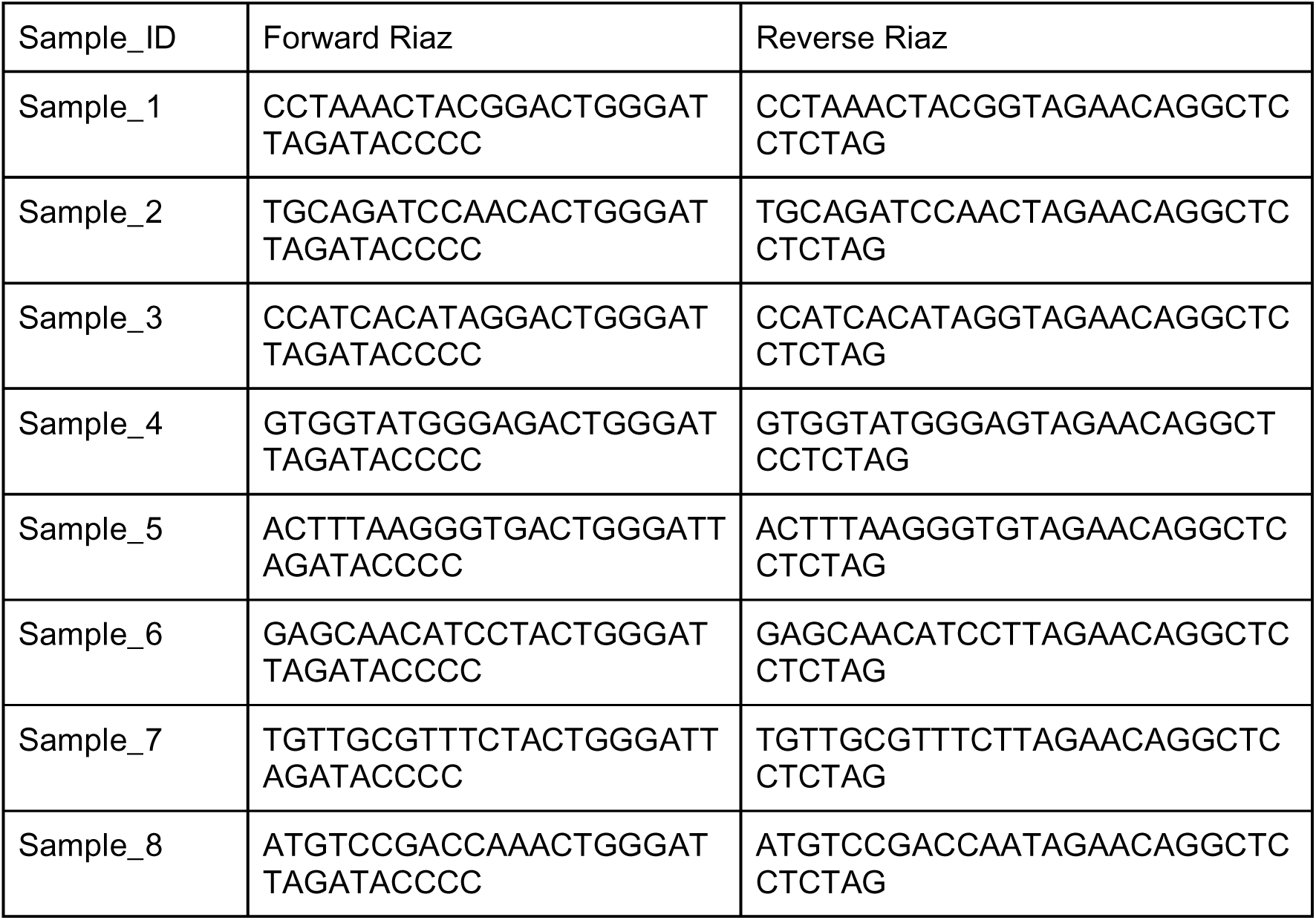
Sequences of the indexed forward and reverse Riaz primers in the 5’-3’ direction.

**Table S4.**
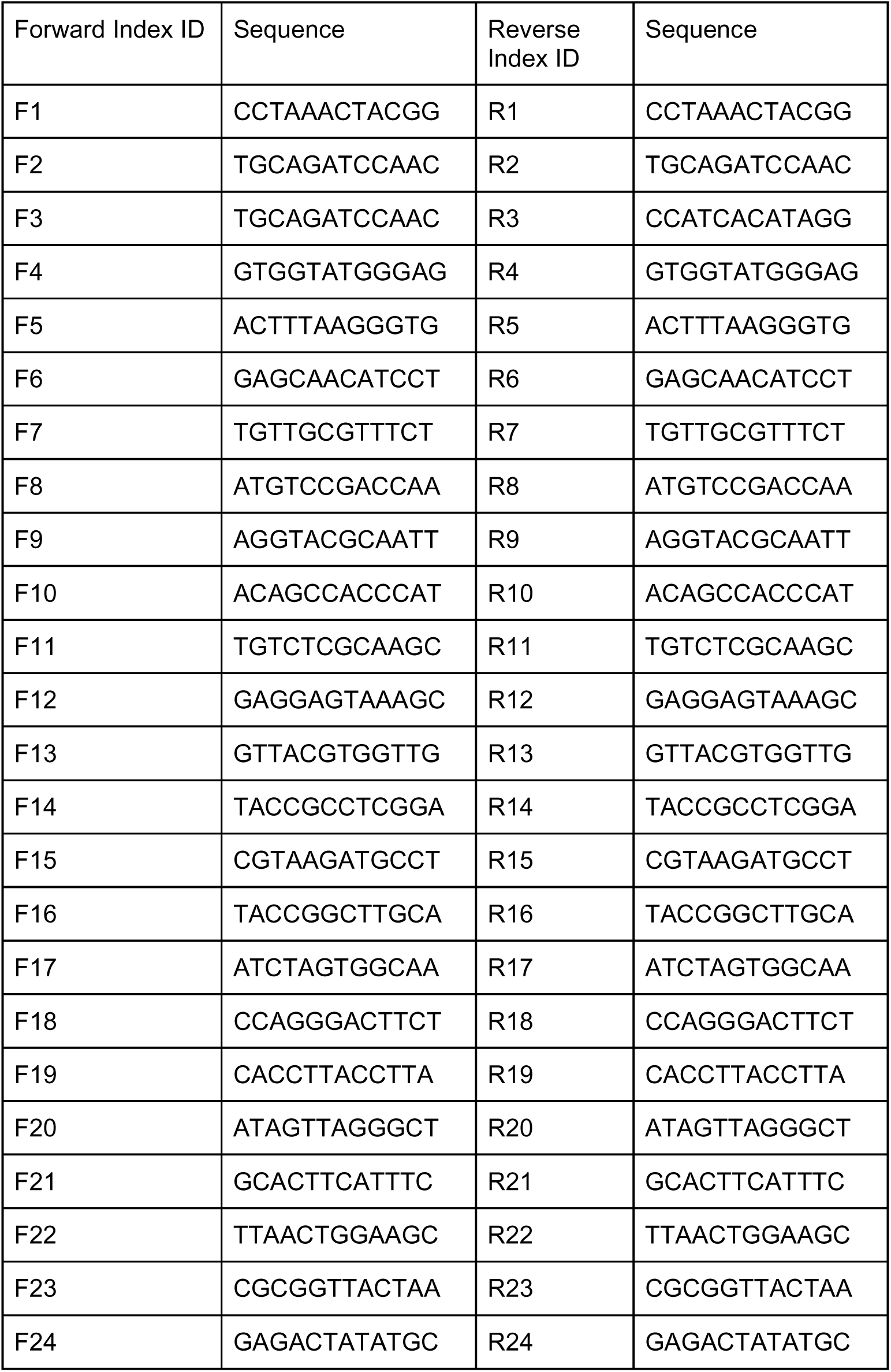
Sequences and IDs of the dual-index set up as described in Fadrosh et al., 2014.

**Figure S1.**
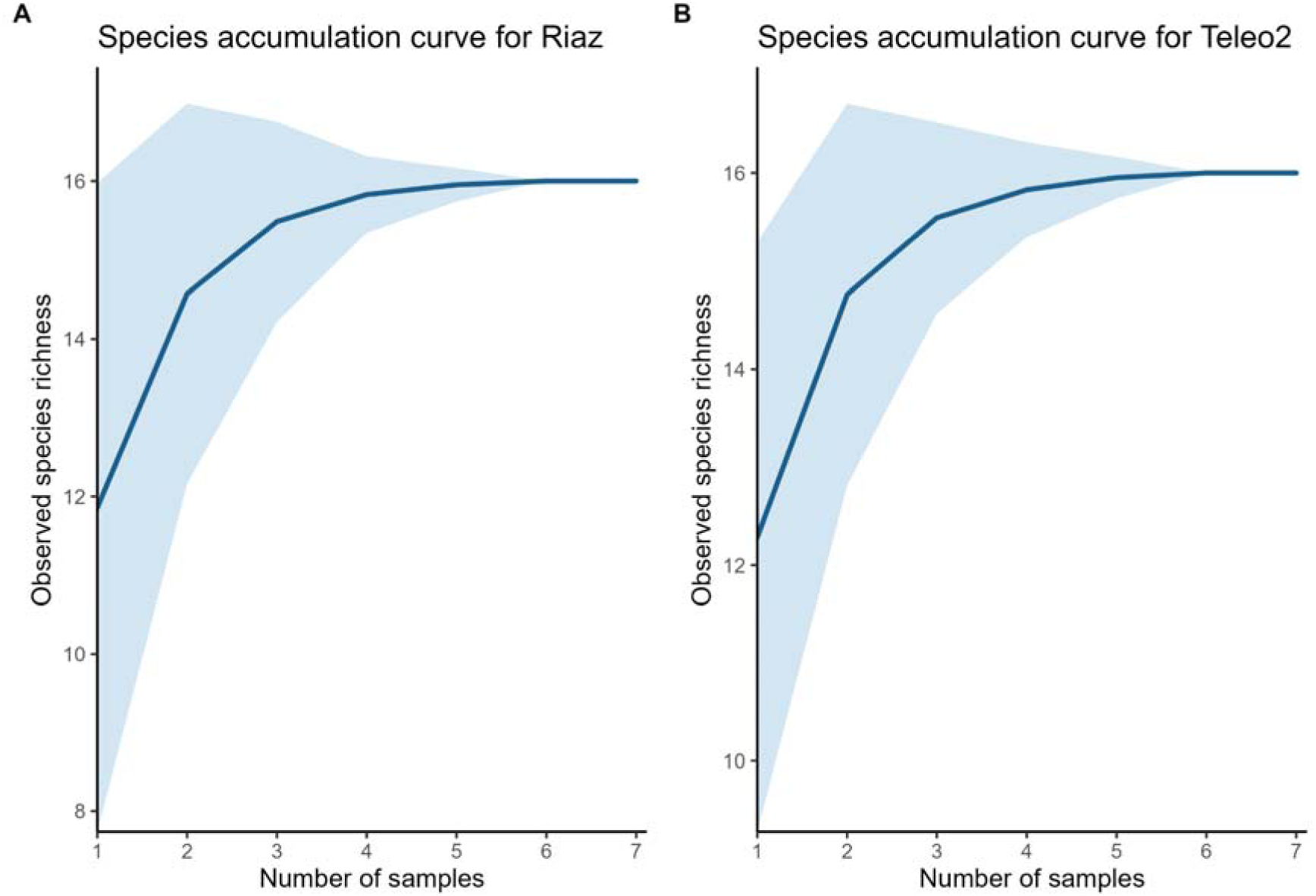
Comparison of species accumulation curves, showing the number of species detected with (A) the Riaz primer set and (B) with Teleo2 accumulating across the 7 samples.

**Figure S2.**
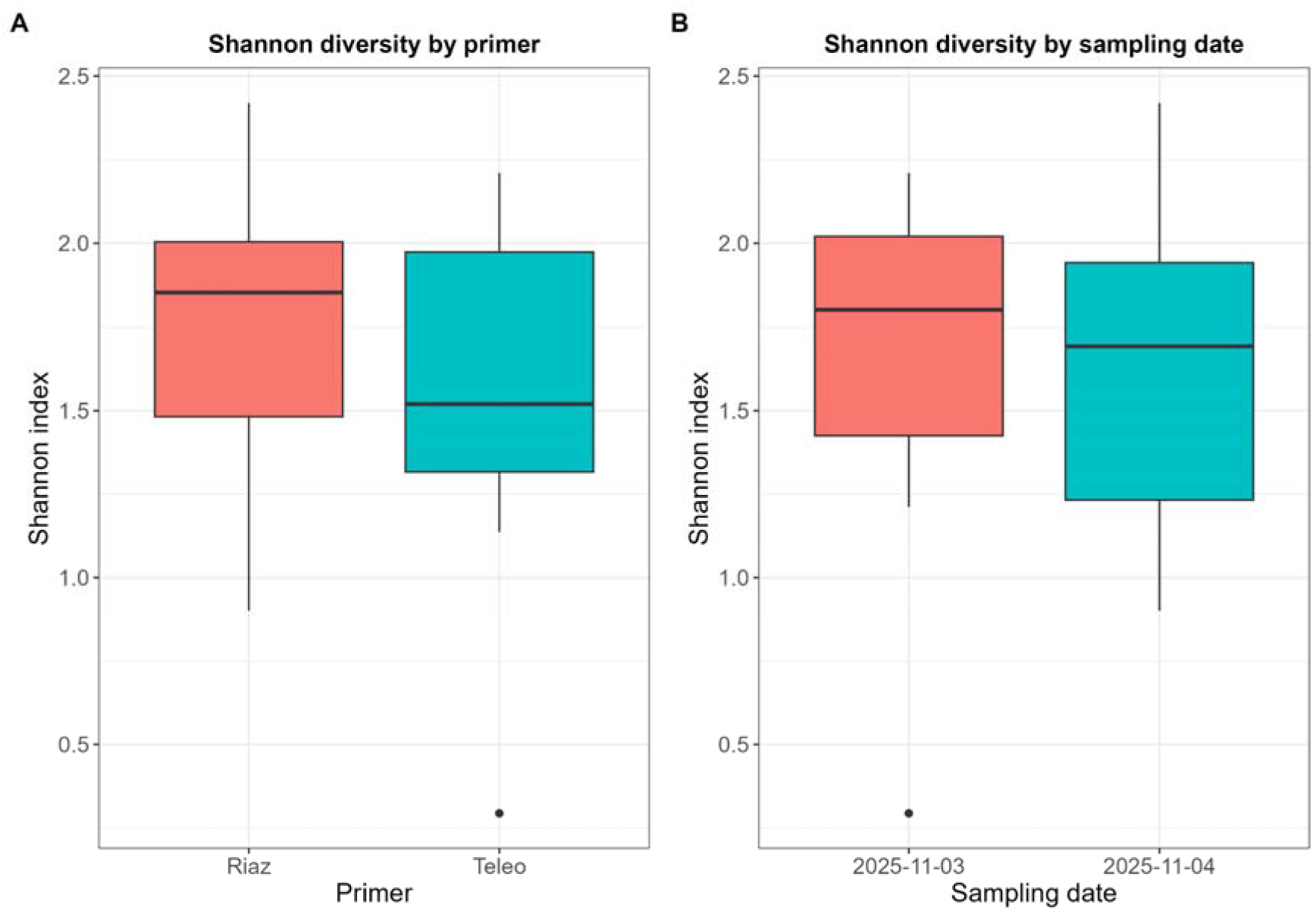
Comparison of (A) the Shannon diversity recovered using the two primer sets Riaz and Teleo2 and (B) comparison of the Shannon diversity recovered on the two different sampling dates.

**Figure S3.**
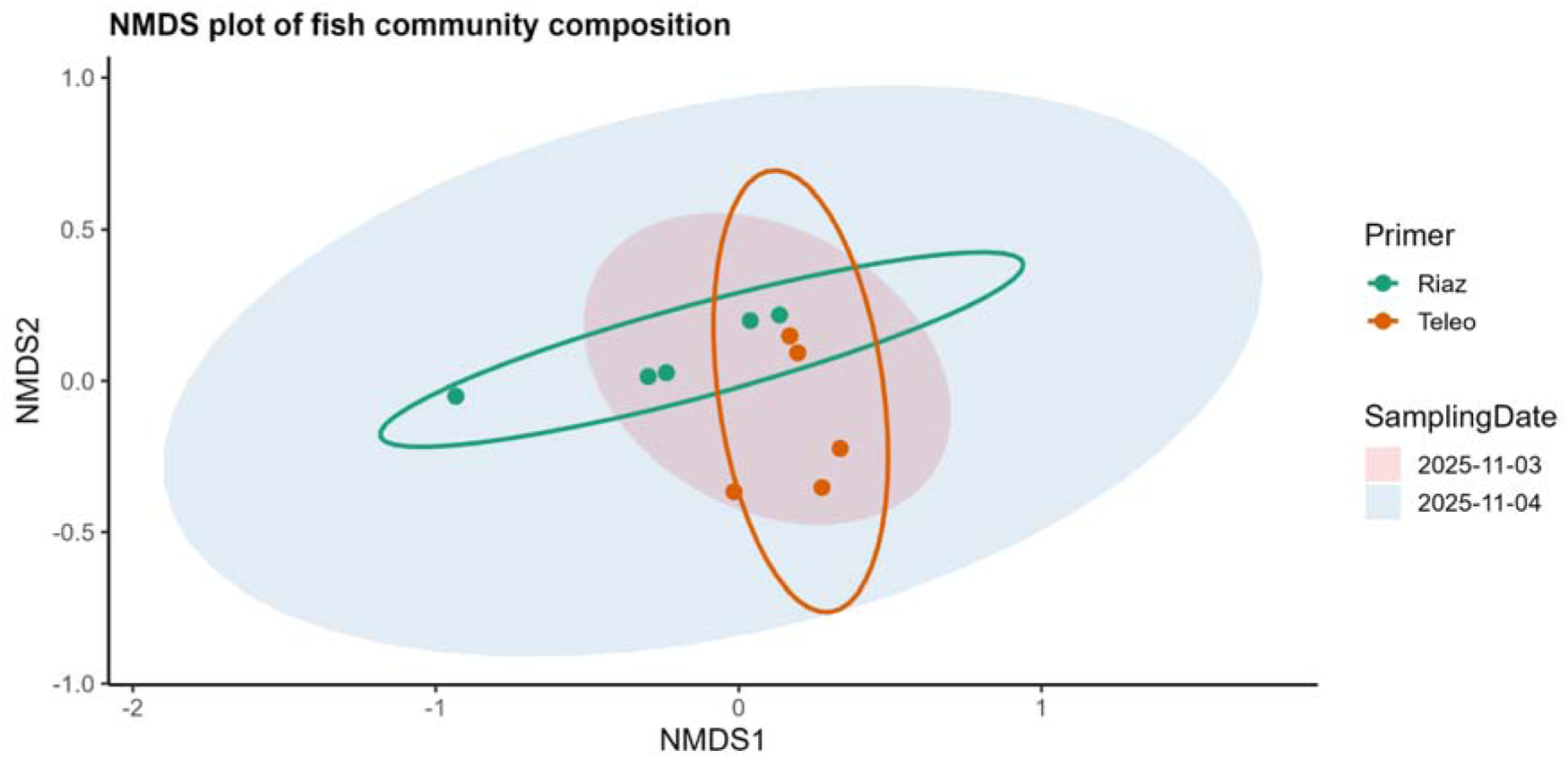
NMDS plot of the fish communities recovered using the two primer sets on the samples collected in the Oslofjord during the two different sampling dates. Due to an identical species composition, Sample 4 (both with Riaz and Teleo2) and Sample 6 (with Teleo2 only) overlap.

**Figure S4.**
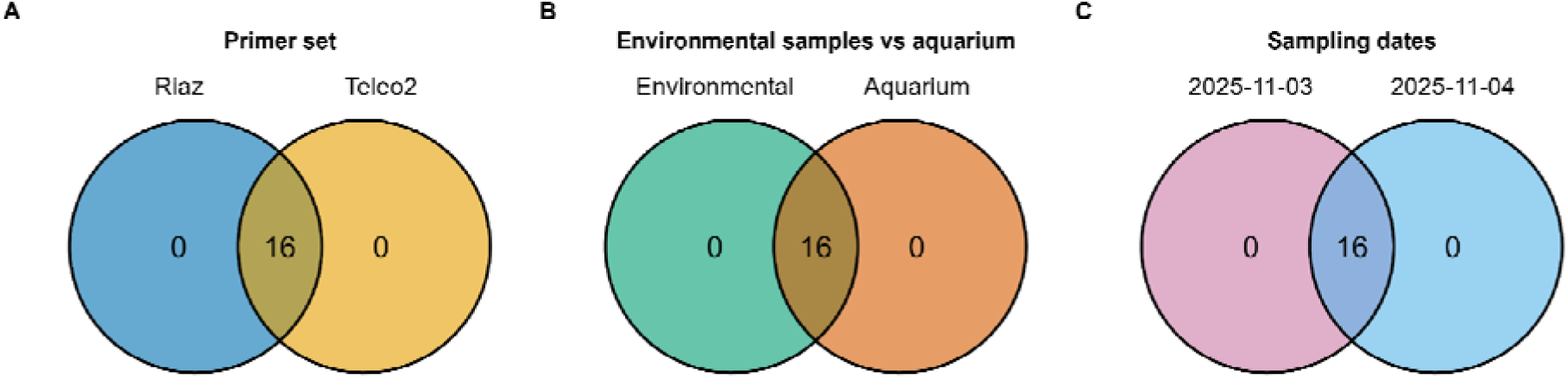
Comparison of the number of species detected by (A) the two primer sets Riaz and Teleo2, (B) in the environmental samples versus in the aquarium samples, and (C) on the two sampling dates.

